# Neutrophil KLF2 regulates inflammasome-dependent neonatal mortality from endotoxemia

**DOI:** 10.1101/2025.02.11.637657

**Authors:** Devashis Mukherjee, Sriram Satyavolu, Asha Thomas, Sarah Cioffi, Yuexin Li, E. Ricky Chan, Katherine Wen, Alex Y. Huang, Mukesh K. Jain, George R. Dubyak, Lalitha Nayak

**Author notes:** Corresponding author: Devashis Mukherjee, Division of Neonatology, UH Rainbow Babies and Children’s Hospital,11100 Euclid Ave, Cleveland, Ohio, USA. Conflicts of interest: All authors declare no competing financial interests.

## Abstract

Preterm neonates die at a significantly higher rate from sepsis than full-term neonates, attributable to their dysregulated immune response. In addition to tissue destruction caused directly by bacterial invasion, an overwhelming cytokine response by the immune cells to bacterial antigens also results in collateral damage. Sepsis leads to decreased gene expression of a critical transcription factor, Krüppel-like factor-2 (KLF2), a tonic repressor of myeloid cell activation. Using a murine model of myeloid-*Klf2* deletion, we show that loss of KLF2 is associated with decreased survival after endotoxemia in a developmentally dependent manner, with increased mortality at postnatal day 4 (P4) compared to P12 pups. This survival is significantly increased by neutrophil depletion. P4 knockout pups have increased pro-inflammatory cytokine levels after endotoxemia compared to P4 controls or P12 pups, with significantly increased levels of IL-1β, a product of the activation of the NLRP3 inflammasome complex. Loss of myeloid-KLF2 at an earlier postnatal age leads to a greater increase in NLRP3 priming and activation and greater IL-1β release by BMNs. Inhibition of NLRP3 inflammasome activation by MCC950 significantly increased survival after endotoxemia in P4 pups. Transcriptomic analysis of bone marrow neutrophils showed that loss of myeloid-KLF2 is associated with gene enrichment of pro-inflammatory pathways in a developmentally dependent manner. These data suggest that targeting KLF2 could be a novel strategy to decrease the pro-inflammatory cytokine storm in neonatal sepsis and improve survival in neonates with sepsis.

**Summary sentence:** KLF2 regulates the developmental response to endotoxin in neonatal mice through the NLRP3 inflammasome signaling pathway.

## INTRODUCTION

Preterm neonates have a high case-fatality rate from bacterial sepsis despite timely antimicrobial therapy, compared to full-term neonates.[1] This has been attributed to dysfunctional innate and adaptive immune systems, early exit from a sterile intra-uterine to a pathogen-laden extrauterine environment, and breaches of physical barriers.

The innate immune system, predominantly myeloid cells, is the first responder against invasive bacterial infections. The preterm immune system is classically labeled as immune suppressed with a decreased ability to mount an inflammatory response. Most studies have investigated whole blood cytokine levels and transcriptomics, mononuclear leukocyte or CD4+ T-cell response, all consistently showing an anti-inflammatory phenotype.[2–5] Neutrophils, the most abundant myeloid cells, are quantitatively and qualitatively distinct in neonates, showing a bone marrow (BM) granulocyte precursor mitotic pool one-tenth the size of adults.[6] Neonates have diminished neutrophil production, creating a smaller storage pool of mature polymorphonuclear cells to be mobilized in response to infection.[6–8] Neonates have decreased neutrophil migratory abilities and reduced transmigration.[9–12] However, neonatal neutrophils demonstrate increased production of pro-inflammatory cytokines compared to adult controls, which can contribute to profound cytokine storm and tissue damage, the hallmark of sepsis-related multi-organ failure and mortality.[13–15] Neonatal neutrophils demonstrate increased interleukin (IL)-1β expression after tumor necrosis factor (TNF)-α and lipopolysaccharide (LPS) stimulation compared to adult neutrophils.[16] In contrast, cord blood monocytes have decreased IL-1β production before 33 weeks gestational age.[17] Here, we identify neutrophils as critical in regulating immune response in preterm neonatal endotoxemia.

IL-1β release in response to microbial pathogens and bacterial cell wall antigens such as LPS is tightly regulated by inflammasome complexes such as the nucleotide-binding oligomerization domain (NOD) leucine-rich repeat (LRR)-, pyrin domain-containing proteins (NLRPs) of which NLRP3 is a prototype.[18] The canonical NLRP3 inflammasome pathway requires two signals. Signal 1 includes extracellular bacterial lipopolysaccharide (LPS), which leads to priming and increased NLRP3 and pro-IL1β transcription and translation directed by the NF-kB transcription factor. Signal 2 includes extracellular ATP, potassium ionophores (nigericin), particulate matter, pathogen-associated RNA, and bacterial and fungal toxins leading to the activation step. Activated NLRP3 inflammasome complex leads to cleavage of pro-caspase-1 into caspase-1, which cleaves pro-IL-1β to mature IL-1β, and gasdermin D (GSDMD) to N-terminal-GSDMD. Aberrant NLRP3 signaling can cause increased cytokine release and is implicated in autoimmune diseases and sepsis – disease states characterized by an inflammatory overdrive. In adult animal sepsis and endotoxemia models, NLRP3 deletion demonstrates protection from sepsis-induced organ injury and shock.[19–22] There are no data on the role of the NLRP3 inflammasome in neonatal neutrophils.[17, 23] We show for the first time that NLRP3 priming is increased in murine BM-derived neutrophils (BMNs) in an age-dependent manner.

Krüppel-like factor-2 (KLF2) belongs to the zinc-finger family of DNA-binding transcription factors and critically regulates the host response to polymicrobial infection and endotoxic shock.[24] Adult mice with myeloid-*Klf2* deletion experience significantly higher mortality than controls after high doses of LPS, attributed to a pro-inflammatory cytokine storm mediated through the HIF-1α pathway.[24] KLF2 regulates neutrophil activation in murine models of heart failure and thrombosis.[25, 26] Pediatric septic shock patients have lower whole-blood KLF2 mRNA levels vs. healthy age-matched controls.[24] There are no data on KLF2 levels in preterm septic neonates or whether KLF2-dependent mechanisms contribute to increased preterm mortality. We show that neutrophil-KLF2 expression is developmentally dependent and regulates NLRP3 expression and activation, potentially contributing to mortality in neonatal pups through NLRP3 signaling.

## METHODS

### Mice

All animal studies were conducted using mice housed in a clean facility and were approved by the Case Western Reserve University (CWRU) Institutional Animal Care and Use Committee. Myeloid-specific *Klf2-*deleted mice were generated in-house by crossing *Klf2*-floxed (*Klf2*^tm2Ling^) with *Lyz2*^Cre^ mice (Jackson Laboratory, ME, USA).[27] *Lyz2*^Cre^ mice express Cre recombinase under the control of the lysozyme-M promoter, specifically expressed in myeloid lineage cells.[28] Myeloid-specific KLF2 transgenic (*Klf2*^Tg^) mice were generated by inserting *Lyz2*^Cre^ mice with a ubiquitin C promoter, followed by a floxed stop codon before the KLF2 gene. Upon recombination, the stop codon is removed, allowing for constitutive expression of *Klf2* in myeloid cells.[27]

a. **Endotoxemia model:** P4 and P12 pups were given intraperitoneal injections of 5 ug/gm LPS (*Escherichia coli* (*E. coli*) strain O55:B5, Sigma Aldrich, MA, USA).
b. **Thermal Imaging:** Thermal images were captured using a FLIR i3 camera (FLIR Systems, Japan). FLIR i3 has a thermal sensitivity of <0.15°C and an accuracy of +/- 2°C and is routinely used for murine thermal imaging.[29, 30] Images were obtained after the injection of LPS and were repeated hourly for up to 8 hours (h), 10cm from the animal. For mice that survived beyond 8h, additional thermal images were taken at 20h, 24h, and 48h.
c. **Neutrophil and macrophage depletion studies:** For neutrophil depletion studies, neonatal mice were injected intraperitoneally with anti-Ly6G, 1A8 clone (BioXCell, NH, USA) at 100 µg/gm. Injections were administered from P1-P3 daily for the P1-3 Ly6G regimen and on P3 only for the P3-Ly6G regimen.[31] Circulating neutrophil depletion efficiency was assessed by flow cytometry. For macrophage depletion, neonatal mice received intraperitoneal injections of 100ug/gm anti-CSF1R antibody (BioXCell NH, USA) daily from P1-P3.[32]
d. **Inhibition of NLRP3 activation:** To inhibit NLRP3 activation, mice were administered MCC950 (Invivogen, CA, USA) reconstituted in phosphate-buffered saline (PBS) daily at 50ug/gm on P3 and P4, 24h apart. Mice received LPS injections 4h after the last dose of MCC950.
e. **Blockade of IL-1 receptor**: To block the binding of IL-1β to the IL-1 receptor, mice were administered commercially available anti-mouse IL-1 receptor antibody (IL-1ra, anti-CD121a, JAMA-147 clone, BioXCell, NH, USA) at a dose of 10ug/gm on P3 and P4, 24h apart.[33] Mice received LPS injections four h after the last dose of IL-1ra.

### Serum extraction

Blood collected from P4 and P12 mice at baseline, four h post-LPS, or sterile PBS injection was allowed to coagulate at room temperature for 30 minutes, centrifuged at 1500g for 15 minutes at 4 °C, and stored at −80°C until further use.

### Multiplex analysis

Serum samples were thawed, diluted, and analyzed using the Mouse Luminex Discovery Assay (R&D Systems; Cat No: LXSAMSM-15) at the CWRU Bioanalyte core facility. This assay allows for multiplex measurement of serum cytokine levels, providing a comprehensive cytokine profile.

### Neutrophil extraction

BMNs were extracted from P4 and P12 pups. Under sterile conditions, bone marrow (BM) cells were collected from bilateral femurs and tibias of each pup at P4 and 12. The bones were crushed in RPMI-1640 media (Gibco, USA), containing 10% fetal bovine serum and penicillin-streptomycin antibiotics (Gibco, USA). Crushed BM was thoroughly mixed and filtered through a 70 µm filter to remove debris. Neutrophils were isolated using the Mouse Neutrophil Enrichment Kit (Stemcell Technologies #19762, Vancouver, BC, Canada). The isolated neutrophils were routinely assessed for purity using flow cytometry (CD45+, CD11b+, Ly6G+), as described previously.[34] The viability of the isolated neutrophils was confirmed by trypan blue staining. At P4 and P12, this method yields ∼ 300,000 and 1.2 million live neutrophils, respectively.

### Inflammasome activation in cell culture

One million neutrophils were plated in 6-well plates in 1mL of RPMI-1640 media supplemented with 2% FBS and primed with LPS 10ng/mL for 2h. They were then treated with nigericin at 20uM for 30 minutes to activate the NLRP3 inflammasome assembly. Cells were kept in a humidified incubator at 37°C in 5% CO_2_. Cells were used for lysate preparation for western blotting, and supernatant was either used for precipitation of proteins for Western blotting or used for ELISA, as described below.

### Enzyme-linked immunosorbent assay (ELISA)

IL-1β ELISA was performed following the manufacturer’s protocol using the Mouse IL-1β ELISA Kit (R&D Systems, #SMLB00C).

### Western blotting

Protein from bone marrow neutrophil lysate was extracted using radioimmunoprecipitation assay (RIPA) buffer supplemented with protease and phosphatase inhibitor cocktails. The protein concentration of the cell lysate was determined using the Pierce BCA Protein Assay Kit (Thermo Fisher Scientific; Cat No:23227). For supernatant, equal amounts (200uL) of supernatant from each arm of the cell culture experiments (1 million BMNs/mL of media) were used for protein precipitation using ProteoExtract Protein Precipitation kit (Sigma Aldrich#539180, USA) using manufacturer-provided protocols. Protein was denatured and reduced by boiling at 95°C for 5 minutes with Laemmli buffer and beta-mercaptoethanol and then separated by polyacrylamide gel electrophoresis. Proteins were then transferred onto polyvinylidene difluoride membranes, blocked for 1 hour at room temperature with 5% bovine serum albumin in PBS, incubated with primary antibodies overnight at 4°C at 1:1000 dilution (NLRP3, Adipogen # AG-20B-0014-C100; IL-1β, R&D Biosciences #AF-401-NA; caspase-1, Adipogen #AG-20B-0042-C100; β-actin, Santa Cruz Biotechnology #sc47778). After washing with PBS-T, horseradish peroxidase-conjugated secondary antibodies were applied at a concentration of 1:5000 for 1 hour at room temperature. Membranes were washed again with PBS-T and developed using Pierce Enhanced Chemiluminescence Western Blotting Substrate (Thermo Scientific #32106) for lysate blots and using SuperSignal West Atto Ultimate Sensitivity Substrate (Thermo Scientific #A38555) for supernatant blots. Protein bands were visualized using a chemiluminescent imager (Azure Biosystems, Dublin, CA, USA).

### Real-time polymerase chain reaction (RT-PCR)

Total RNA was extracted from BMNs (RNeasy Micro Kit, QIAGEN) and reverse transcribed to complementary DNA using the iScript Reverse Transcription Kit (Bio-Rad, 170-8841). RT-PCR was performed using the Universal SYBR Green method on the StepOnePlus Real-Time PCR System (Applied Biosystems). Relative gene expression of NLRP3 was calculated using the ΔΔCt method, normalized to the expression of RPL13A (ribosomal protein L13a), a housekeeping gene used for internal control. Primer sequences are detailed in Table S1.

### Lactate dehydrogenase (LDH) Assay

To assess cellular cytotoxicity by LDH release, the CyQuant LDH Cytotoxicity Assay (Invitrogen by Thermo Fisher Scientific; Cat No C20300) kit was used per the manufacturer’s protocols. BM-derived neutrophils were seeded in triplicate at 10,000 cells per well in a 96-well plate and subjected to LPS 10ng/mL for 2 hours for priming, followed by nigericin 20uM for NLRP3 inflammasome activation. An equal number of cells were seeded simultaneously and treated with a company-provided lysis buffer to assess the maximum LDH activity. Cells in media without LPS acted as controls for spontaneous LDH activity. Absorbances were measured at 490nm and 680nm. To calculate LDH activity, the background activity (680nm) was subtracted from the 490nm absorbance. The spontaneous LDH activity was subtracted from the maximum LDH activity as well as the LDH activity of the LPS/nigericin-treated cells. The percentage of cytotoxicity was calculated as a percentage of LDH activity in LPS/nigericin-treated cells as a fraction of the maximum LDH activity. A manufacturer-provided LDH positive control was used for all assays.

### RNA-sequencing and analysis

Total RNA from P4 and P12 pups (n=7 from each genotype) was used for library preparation. RNA was extracted from BM neutrophils as described above. At P4, RNA from two animals was pooled to meet the minimum RNA quantity for sequencing. Quality control of total RNA samples was performed using a Qubit Fluorometer (Invitrogen) for RNA quantification and Fragment Analyzer 5200 (Agilent) to assess RNA quality using a cut-off of RIN > 7.0 to select specimens for further analysis. Ribosomal RNA was depleted (NEBNext rRNA Depletion Kit v2 (Human/Mouse/Rat, New England Biolabs, USA), and this rRNA-depleted RNA was used as input for the NEBNext Ultra II Directional RNA Library Prep kit for Illumina (New England BioLabs, USA) in which libraries were tagged with unique adapter-indexes. Final libraries were validated on the Fragment Analyzer, quantified via qPCR, and pooled at equimolar ratios. Pooled libraries were diluted, denatured, and loaded onto the Illumina NovaSeq X sequencing system, following the Illumina User Guide for a paired-end run. Sequencing reads generated from the Illumina platform were assessed for quality and trimmed for adapter sequences using TrimGalore! v0.4.2 (Babraham Bioinformatics), a wrapper script for FastQC and cutadapt. Reads that passed quality control were aligned to the mouse reference genome (mm10) using the STAR aligner v2.5.3.[35] The aligned reads were analyzed for differential expression using Cufflinks v2.2.1. This RNAseq analysis package reports the fragments per kilobase of exon per million fragments mapped (FPKM) for each gene using annotations from GENCODE annotation for mm10.[36] Differential analysis was performed using the cuffdiff command in a pairwise manner for each group. Differential genes were identified using a significance cutoff of q-value < 0.05. The genes were subjected to gene set enrichment analysis (GenePattern, Broad Institute) and cross-referenced using the Gene Ontology database to determine any relevant processes that may be differentially over-represented for the conditions tested. Additional pathway analysis was performed using iPathwayGuide (Advaita Bio). Meta-analysis and gene ontology results were generated using iPathwayGuide’s built-in software.

### Statistical Analysis

GraphPad Prism was used for all statistical analysis except for RNA-sequencing, which is detailed above. Chi-square tests were used to analyze survival benefit, with a p-value of 0.05 considered significant. Western blots were quantified as optimal densities using ImageJ (National Institutes of Health, Bethesda, MD, USA) and normalized to beta-actin, a standard housekeeping gene. For supernatant blots, equal volumes of supernatant from equal cell seeding densities were used for gel electrophoresis to ensure equal protein loading amounts. Two-way ANOVA was used for Western blots, ELISA, and LDH assay results to analyze the effect of treatment (control vs. LPS+nigericin) and loss of KLF2 (*Klf2*^fl/fl^ *Lyz2*^Cre^ vs. *Lyz2*^Cre^) on protein expression and measured levels.

## RESULTS

### Neutrophil KLF2 regulates LPS-induced mortality in an age-dependent manner

To investigate whether decreased myeloid-*Klf2* gene expression is responsible for the increased mortality in preterm neonates, we used a myeloid-specific *Klf2* knockout murine model of endotoxemia. LPS (5ug/gm) was injected at P4 and P12 to reflect preterm and term innate immune responses, respectively (Fig. 1A).[37] At P4, *Klf2*^fl/fl^*Lyz2*^Cre^ pups experienced significantly decreased survival than *Lyz2*^Cre^ pups (20%, n=40 vs. 59%, n=39, p=0.0004, Fig. 1C). At P12, there is 100% survival in both groups (n=20, Fig. 1C) with a 5ug/gm LPS dose. P4 pups also experience a significantly greater degree of hypothermia vs. P12 pups (Fig. 1D, E), regardless of genotype. Although P4 *Klf2*^fl/fl^*Lyz2*^Cre^ had a decreasing trend in their core body temperatures compared to P4 *Lyz2*^Cre^ pups, it did not reach statistical significance before the onset of mortality in these pups.

**Figure 1.**
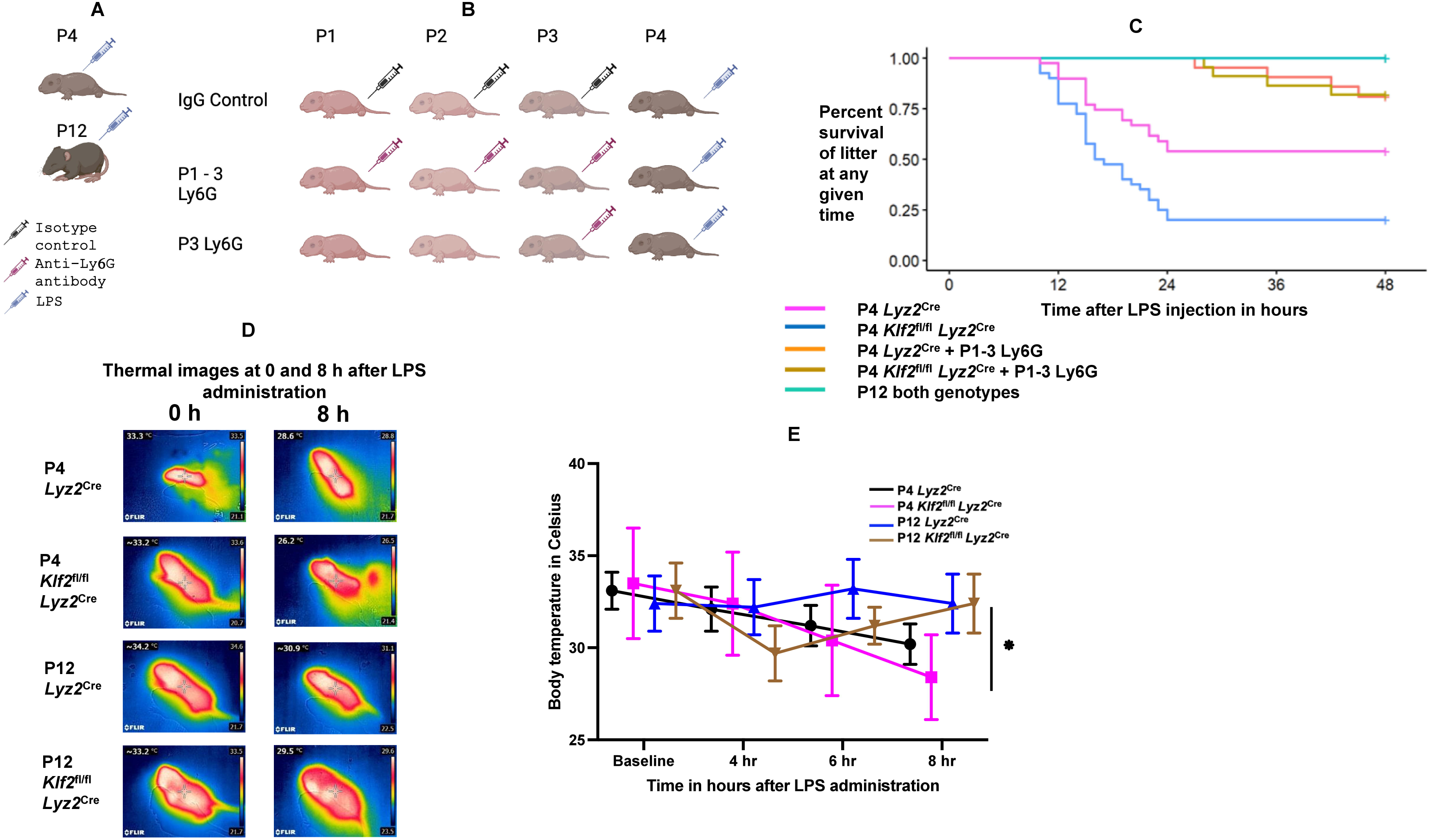
Loss of myeloid-KLF2 leads to increased mortality at an earlier postnatal age through neutrophils. (A) Postnatal day (P) 4 and P12 pups were injected with 5ug/gm of E. coli O55:B5 LPS intraperitoneally. (B) Neutrophil depletion was performed in P4 pups using anti-Ly6G antibody injections either from P1-3 (P1-3 Ly6G) or only on P3 (P3 Ly6G), followed by LPS 24h after the last antibody injection. Rat IgG isotype controls were used for control experiments. (C) P12 pups (both *Klf2*^fl/fl^*Lyz2*^Cre^ and *Lyz2*^Cre^, n=20 each group) have 100% survival 48h after LPS injection, whereas at P4, *Klf2*^fl/fl^ *Lyz2*^Cre^ experience 81% mortality within 48h (n=40) and *Lyz2*^Cre^ pups have 40% mortality (n=39) within 48h. P3 Ly6G decreases mortality at P4 for both genotypes, with survival increasing to 79% in *Klf2*^fl/fl^*Lyz2*^Cre^ and 78% in *Lyz2*^Cre^ pups. (D) and (E) show thermal images and temperature changes after LPS administration, respectively. P4 pups experience increased hypothermia compared to P12, and the loss of myeloid-KLF2 exacerbates this hypothermia. Images were obtained using a FLIR i3 camera at a 10cm from each pup, at baseline, and then hourly. Temperature graphs are not shown beyond the 8-hour time point to reduce the introduction of selection bias in the differences between temperatures as pups started dying after the 8-hour time point.

Since *Lyz2^Cre^* is expressed in both neutrophils and monocytes/macrophages, we performed separate neutrophil and macrophage depletion studies to determine the cell-specific contribution of decreased KLF2 to the increased mortality. We have shown previously that in C57Bl/6 pups (Fig. 1B), anti-Ly6G antibody on P3 (P3-Ly6G) reduces circulating neutrophil count by 50%, and three days of daily anti-Ly6G antibody administrations from P1–P3 (P1-3 Ly6G) reduces it by 73%.[31] P3-Ly6G (50% neutrophil depletion) followed by LPS on P4 significantly increased survival in *Klf2*^fl/fl^*Lyz2*^Cre^ pups (20% to 50%, Fig. S1), but the increase in survival seen in *Lyz2*^Cre^ pups was not significant (59% to 65%, Fig. S1). P1-3 Ly6G (73% neutrophil depletion) followed by LPS on P4 however led to a significant increase in survival in both *Klf2*^fl/fl^*Lyz2*^Cre^ (20% to 79%) and in *Lyz2*^Cre^ (59% to 78%, Fig. 1C, Fig. S1) pups. In contrast, macrophage depletion (anti-CSF1R antibodies) did not reduce mortality in either genotype (data not shown). These data demonstrate that neutrophil-KLF2 is critical in regulating the age-dependent mortality from endotoxemia and that survival after endotoxemia is neutrophil-dependent.

We then investigated whether constitutive myeloid *Klf2* overexpression from birth in the myeloid cells decreases mortality after endotoxemia. *Klf2*^Tg^ pups received 5 ug/gm LPS i.p. on P4 and were followed closely for the next 48 hours. Five out of 19 pups (3 separate litters) experienced lethality, indicating that while *Klf2*^Tg^ pups have significantly less mortality after LPS at P4 compared to *Klf2*^fl/fl^ *Lyz2*^Cre^ pups (41% vs. 80%, p<0.0001, Fig. S1), *Klf2* overexpression does not confer a statistically significant survival benefit over the control *Lyz2*^Cre^ genotype (p=0.27, Fig. S1).

### Myeloid-KLF2 regulates the age-dependent pro-inflammatory response

To understand the contribution of different cytokines to the increased mortality seen at an earlier postnatal age, we performed a multiplex analysis of pro- and anti-inflammatory cytokines. Sera were collected 4h after PBS or LPS exposure (Fig. 2). We found an exaggerated pro-inflammatory response in *Klf2*^fl/fl^*Lyz2*^Cre^ vs. *Lyz2*^Cre^ pups. Further, this is significantly different at P4 vs. P12. IL-1β was the most significantly and consistently elevated cytokine in P4 *Klf2*^fl/fl^*Lyz2*^Cre^ pups, which also had higher levels of most pro-inflammatory cytokines, including IL-1β, IFN-γ, TNF-α, and CCL3 (MIP-1α). Anti-inflammatory cytokines such as IL-4 were decreased in P4 *Klf2*^fl/fl^*Lyz2*^Cre^ pups compared to P4 *Lyz2*^Cre^ pups. IL-10 was the only anti-inflammatory cytokine significantly increased in P4 *Klf2*^fl/fl^*Lyz2*^Cre^ pups. These data show that myeloid-*Klf2* deletion is associated with an exaggerated pro-inflammatory response to sterile inflammation. Further, this response is age-dependent and more pronounced at an earlier neonatal age.

**Figure 2.**
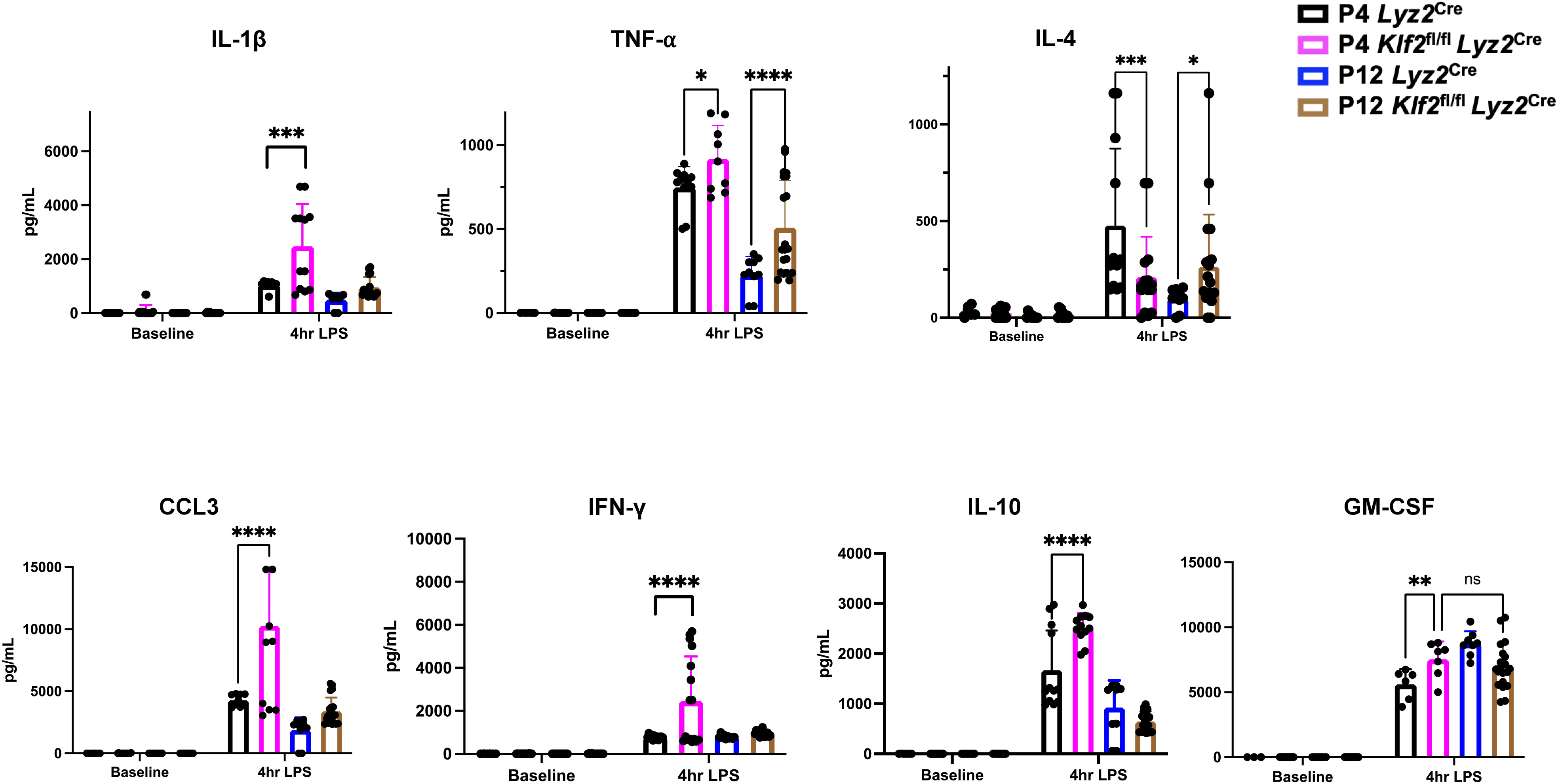
Multiplex analysis of pro- and anti-inflammatory cytokines on serum obtained four h after PBS or LPS treatment in P4 and P12 *Klf2*^fl/fl^*Lyz2*^Cre^ and *Lyz2*^Cre^ pups. Data are expressed as pg/mL for all cytokines. Each sample was run in duplicate and averaged to obtain a single data point.

### Neutrophil-KLF2 affects NLRP3 inflammasome priming

IL-1β is a critical pro-inflammatory cytokine significantly elevated in P4 *Klf2*^fl/fl^*Lyz2*^Cre^ pups. IL-1β is primarily released from priming and activation of the NLRP3 inflammasome. The priming step, triggered by extracellular LPS, leads to increased *Nlrp3* gene expression and is under the transcriptional control of NF-kB.[18] Under physiologic conditions, KLF2 inhibits the recruitment of NF-kB co-activators to its promoter sites.[24, 38] We hypothesized that loss of KLF2 leads to increased NLRP3 signaling through increased priming or activation, thereby increasing IL-1β release. Since we identified neutrophils as critical regulators of mortality in our model, we focused on this specific cell for our studies.

P4 and P12 *Klf2*^fl/fl^*Lyz2*^Cre^ and *Lyz2*^Cre^ BMNs were LPS-primed and treated with nigericin, a potassium ionophore that activates NLRP3 inflammasome assembly. *Klf2*^fl/fl^*Lyz2*^Cre^ neutrophils demonstrate significantly higher NLRP3 protein expression both at baseline and after LPS priming than age-matched *Lyz2*^Cre^ neutrophils (Fig. 3A and B). Consistently, cleavage of pro-IL-1β (evidenced by decreased expression of pro-IL-1β in the whole cell lysate) and release of mature IL-1β into the supernatant (evidenced by increased mature IL-1β in the supernatant western blots and by supernatant IL-1β ELISA) is significantly greater in activated *Klf2*^fl/fl^*Lyz2*^Cre^ vs. *Lyz2*^Cre^ neutrophils (Fig. 3C, D, E), and this was enhanced at P4 vs. P12. Activation of the NLRP3 inflammasome assembly after the priming step leads to the cleavage of pro-caspase-1 into caspase-1 p20, which is necessary for the proteolytic cleavage of pro-IL-1β into mature IL-1β. Since activated P4 *Klf2*^fl/fl^ *Lyz2*^Cre^ BMNs have increased release of mature IL-1β into the supernatant, we then measured pro-caspase-1 cleavage. Release of caspase-1 p20 into the supernatant was significantly higher in *Klf2*^fl/fl^ *Lyz2*^Cre^ vs. *Lyz2*^Cre^ BMNs, which was enhanced at P4 vs. P12 (Fig. 3C, D). Although there was a trend towards increased pro-caspase-1 cleavage in the whole cell lysate in P4 *Klf2*^fl/fl^ *Lyz2*^Cre^ BMNs, this was not statistically significant (Fig. 3C, D) Interestingly, at P4, even in the absence of priming and activation stimuli, *Klf2*^fl/fl^ *Lyz2*^Cre^ BMNs demonstrate the presence of caspase-1 p20 in the supernatant. No differences were observed in the LDH release between the four groups (Fig. S2), suggesting that the increase in mature IL-1β release by *Klf2*^fl/fl^*Lyz2*^Cre^ vs. *Lyz2*^Cre^ neutrophils was non-pyroptotic. Although NLRP3 protein expression was higher in *Klf2*^fl/fl^*Lyz2*^Cre^ vs. *Lyz2*^Cre^ neutrophils, this was not observed at a transcriptional level at either age (Fig. S3). Together, these results show that KLF2 non-transcriptionally and developmentally regulates neutrophil-NLRP3 priming and activation.

**Figure 3.**
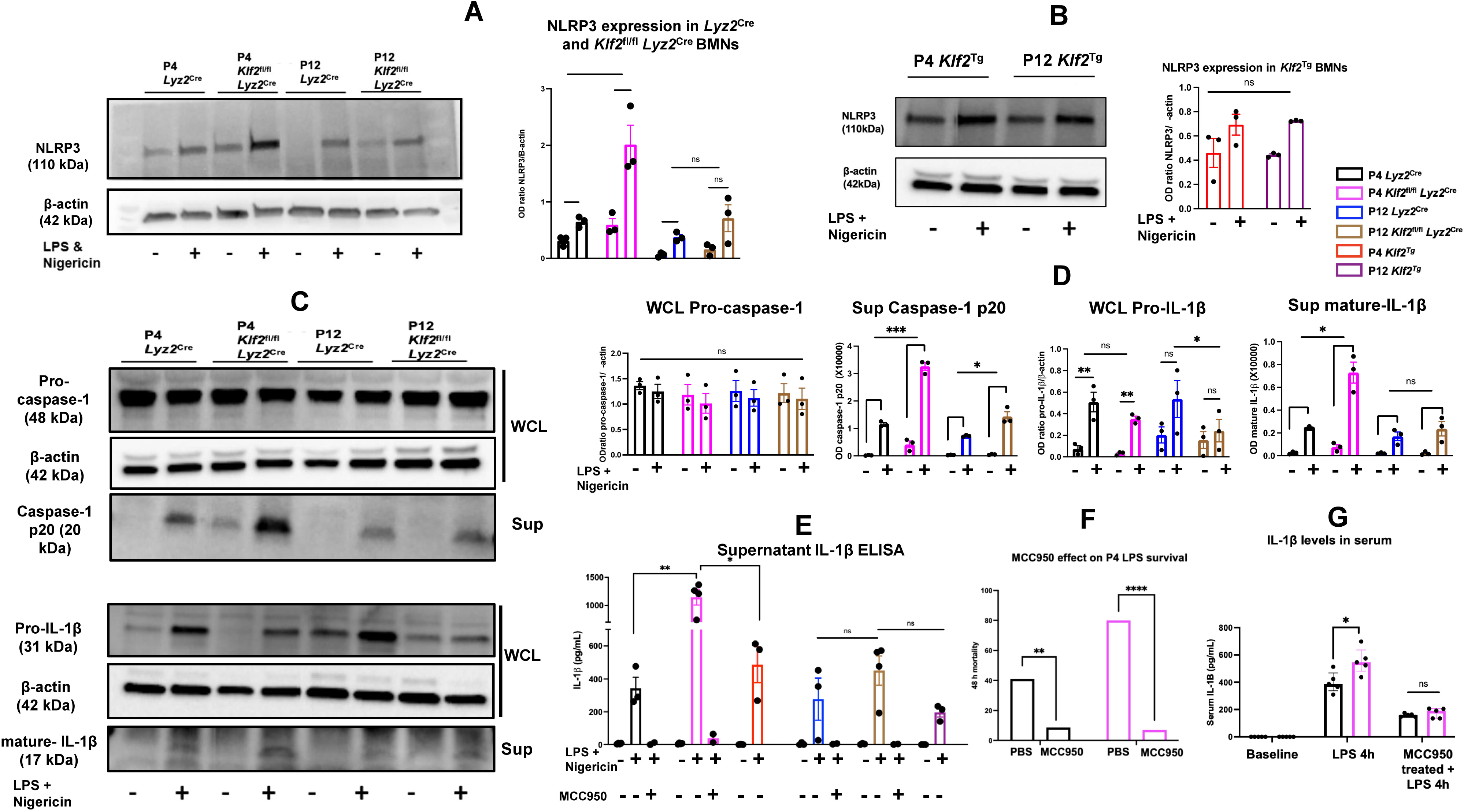
KLF2 regulates NLRP3 protein expression in neonatal neutrophils. (A) shows protein expression of NLRP3 in the whole cell lysate of bone marrow-derived neutrophils primed with 10ng/mL LPS for two h followed by nigericin stimulation at 20uM for 30 minutes to activate the NLRP3 inflammasome. 20ug of protein extracted from the cell lysate was loaded per well and separated using SDS-PAGE. NLRP3 expression was normalized to beta-actin. For P4, 6-7 pups were pooled to obtain a n of 1; for P12, 2-3 pups were pooled to obtain a n of 1. KLF2 deletion increases NLRP3 protein expression in neutrophils, both in unprimed and LPS-primed cells. This change is more evident at P4, with P4 *Lyz2*^Cre^ neutrophils having increased NLRP3 protein expression than P12 *Lyz2*^Cre^ neutrophils. (B) shows NLRP3 protein expression in BMNs obtained from P4 and P12 *Klf2*^Tg^ pups, with significant increase in NLRP3 expression on priming, but no difference in the increase in expression between the two postnatal ages. (C) BMNs from P4 and P12 *Klf2*^fl/fl^ *Lyz2*^Cre^ pups were plated at a concentration of 1 million cells/mL in RPMI-1640 and treated with LPS at 10ng/uL for 2 hours, followed by nigericin at 20uM for 30 mins. Whole cell lysates were prepared for gel electrophoresis, and simultaneously equal volumes of supernatant were precipitated (to ensure normalization of the secreted proteins in the supernatant) to extract for protein for 1-D gel electrophoresis. Blots were probed for pro-IL-1β, pro-caspase-1, and β-actin in the whole cell lysate, and for mature-IL-1β and caspase-1 p20 in the supernatant. Although pro-caspase-1 expression did not differ significantly between the ages and genotypes with LPS and nigericin treatment, the caspase-1 p20 levels in the supernatant were significantly higher in activated *Klf2*^fl/fl^ *Lyz2*^Cre^ BMNs compared to *Lyz2*^Cre^ BMNs at P4 (p=0.0007, two-way ANOVA, and this difference was less pronounced at P12 (p=0.0198, two-way ANOVA). P4 *Klf2*^fl/fl^ *Lyz2*^Cre^ BMNs have increased caspase-1 p20 in their supernatant even without LPS priming or nigericin activation. Pro-IL-1β cleavage is increased after LPS and nigericin treatment with decreased pro-IL-1β expression in the whole cell lysate and increased mature IL-1β in the supernatant. Mature IL-1β levels were significantly higher in the supernatant of LPS-primed *Klf2*^fl/fl^*Lyz2*^Cre^ vs. *Lyz2*^Cre^ BMNs at P4 (p=0.0105, two-way ANOVA) but not at P12 (p=0.0524), mirroring the ELISA results in (E), which also demonstrates that MCC950 treatment of neutrophils decreases IL-1β release by *Klf2*^fl/fl^*Lyz2*^Cre^ and *Lyz2*^Cre^ neutrophils at P4 and P12 (n=2 each). (F) MCC950 administration in vivo decreases mortality in P4 *Klf2*^fl/fl^*Lyz2*^Cre^ and *Lyz2*^Cre^ pups. P3-4 MCC950 – 50ug/gm on P3 and P4, followed by LPS 4 h later, significantly reduced the 48-hr mortality to 7% (n=21) from a baseline of 81% in *Klf2*^fl/fl^ *Lyz2*^Cre^ pups (p<0.00001), and in P4 *Lyz2*^Cre^ pups from 81% to 8.6% (n=35, p=0.0014) (G) Serum obtained from mice treated with P3-4 MCC950 or PBS, followed by LPS at P4, and then harvested at 4 h post LPS showed a significant decrease in IL-1β levels by ELISA. The IL-1β levels in MCC950-treated *Klf2*^fl/fl^*Lyz2*^Cre^ and *Lyz2*^Cre^ pups were not significantly different after LPS administration, but they were in the PBS-treated pups.

We then measured NLRP3 priming and activation in BMNs isolated from P4 and P12 *Klf2*^Tg^ pups to identify whether constitutive myeloid *Klf2* gene overexpression leads to the abolition of the postnatal age-dependent change in NLRP3 protein expression and IL-1β release in BMNs. LPS-priming leads to increased NLRP3 expression as expected in *Klf2*^Tg^ BMNs at both P4 and P12 (Fig 3B), but there was no difference in NLRP3 protein expression between P4 and P12 BMNs, unlike in *Klf2*^fl/fl^ *Lyz2*^Cre^ and *Lyz2*^Cre^ BMNs, where there is a strong postnatal age-dependent change in NLRP3 protein expression (Fig. 3A, B). In addition, NLRP3 protein expression in stimulated P4 BMNs was significantly lower in BMNs from *Klf2*^Tg^ pups vs. in BMNs from *Klf2*^fl/fl^ *Lyz2*^Cre^ pups (Fig. S4). However, it was not lower than those seen in BMN from *Lyz2*^Cre^ pups. These differences in BMN NLRP3 expression levels correlated with the differences in LPS-mortality results seen across all three strains. IL-1β release into the supernatant was also not significantly different between P4 and P12 BMNs in from the *Klf2*^Tg^ pups, and was lower in stimulated P4 *Klf2*^Tg^ BMNs compared to stimulated *Klf2*^fl/fl^ *Lyz2*^Cre^ BMNs (p=0.0173), but not when compared to P4 *Lyz2*^Cre^ BMNs(Fig. 3E).

### Postnatal day – 4 mortality is dependent on NLRP3 inflammasome activation

We then postulated that LPS-induced mortality in P4 pups is mediated through NLRP3-inflammasome signaling. MCC950 specifically inhibits NLRP3 activation without affecting other inflammasomes.[39] We administered 50ug/gm MCC950 24h before, followed by a second dose 4 hours before the LPS injection. MCC950 significantly decreased mortality in P4 *Klf2*^fl/fl^*Lyz2*^Cre^ pups (80% to 7%, n=21, Fig. 3F). This regimen also reduced mortality in *Lyz2*^Cre^ pups to 8.6% (n=35, Fig. 3F). Increasing the dose of MCC950 beyond 50ug/gm did not confer additional survival benefit. Consistently, sera obtained from pups 4h after LPS administration in MCC950-treated animals showed decreased levels of IL-1β compared to PBS-treated pups (Fig. 3G). Although IL-1β levels decreased to similar levels in MCC950-treated *Klf2*^fl/fl^*Lyz2*^Cre^ and *Lyz2*^Cre^ pups after LPS, the decrease in IL-1β was more significant in *Klf2*^fl/fl^*Lyz2*^Cre^ vs. *Lyz2*^Cre^ Cre pups at P4. To study the effect of MCC950 specifically on BMNs, these cells were treated ex vivo with either MCC950 or DMSO followed by LPS and nigericin to prime and activate the NLRP3 inflammasome. MCC950 decreases IL-1β release significantly in all four arms compared to DMSO-treated neutrophils, with the difference between the IL-1β decrease in *Klf2*^fl/fl^*Lyz2*^Cre^ and in *Lyz2*^Cre^ BMNs being significant (Fig. 3E). Thus, the increase in IL-1β seen in neutrophils from the loss of KLF2 is driven through the NLRP3 inflammasome pathway. To identify whether inhibition of the IL-1 receptor would decrease mortality after endotoxemia, we administered 10ug/gm of IL-1 receptor antagonist (IL-1ra) on P3 and P4, followed by LPS injection 4 hours after the last dose of IL-1ra, mimicking the MCC950 dosing schedule. IL-1r blockade led to 100% survival after LPS in P4 *Klf2*^fl/fl^ *Lyz2*^Cre^ pups and 93.8% in P4 *Lyz2*^Cre^ pups (Fig. S5).

These results confirm that LPS-induced mortality in neonatal mice is dependent on NLRP3-inflammasome assembly and the subsequent production of IL-1β. While we cannot conclusively identify the cellular determinants of the survival benefit noted with NLRP3 inhibition, our ex vivo studies showing heightened NLRP3 activation in the *Klf2*-deficient neutrophils suggest that the neutrophil-specific NLRP3 activity must contribute to the overall mortality noted in our animal model.

### KLF2 regulates age-associated alterations in the neutrophil transcriptome

We then performed transcriptomic analysis of P4 and P12 *Klf2*^fl/fl^*Lyz2*^Cre^ and *Lyz2*^Cre^ BMNs to investigate how the loss of a critical transcription factor, KLF2 alters the neutrophil transcriptomic profile developmentally, and whether this contributes to the skewed pro-inflammatory profile. Principal component analysis showed a clear separation between the four groups (Fig. 4A) and was stronger at P12 than at P4. *Klf2* gene expression was not different between P4 and P12 *Lyz2*^Cre^ BMNs but was lower in P4 vs. P12 *Klf2*^fl/fl^ *Lyz2*^Cre^ BMNs (Fig. S6). At P4 and P12, there were 304 and 661 significantly (threshold p-value 0.05) differentially expressed genes (DEGs) between *Klf2*^fl/fl^*Lyz2*^Cre^ and *Lyz2*^Cre^ pups, respectively (Fig. 4 B). The top biological processes impacted at both P4 and P12 by myeloid-*Klf2* deletion involved immune response to bacteria, innate immune regulation, and cellular response to chemokines (Fig. 4C). At P4, the significantly different hallmark response pathways (FDR <0.05) were inflammatory response (14 genes), IFNγ response (13 genes), myogenesis (11 genes), and TNF-α signaling through NF-kB (10 genes). This differs at P12, with seven significantly different hallmark response pathways (Table S2). Inflammatory response and TNF-signaling through NF-kB were the only two significantly different pathways between *Klf2*^fl/fl^*Lyz2*^Cre^ and *Lyz2*^Cre^ at both P4 and P12. At P12, these pathways included the downregulated DEGs *Hbegf*, *Ccrl2*, *Ldlr, Ripk2, Il7r,* and *Nampt*, and the upregulated DEGs, *Ccl2, Sphk1, Pde4b*, and *Btg2.* At P4, these pathways included only downregulated DEGs *Ccrl2, Ccl2,* and *Ldlr*. We then combined all significant DEGs at P4 and P12 to create a meta-analysis to identify key biological pathways in neutrophils responsible for the developmentally regulated phenotype seen with myeloid-KLF2 deletion. This included 15 biological processes, out of which those that involved the most DEGs both at P4 and P12 were negative regulation of molecular function, immune response-regulating cell surface receptor signaling pathway, type II interferon production regulation, and leukocyte-mediated cytotoxicity. Within the meta-analysis, there were 14 pathways, out of which the NF-kB signaling pathway and Toll-like receptor signaling pathway were both enriched (Fig. 4D). It is important to note that both of these signaling pathways are key upstream regulatory pathways controlling NLRP3 inflammasome and interleukin signaling. Out of the top 25 upstream regulatory genes at P4 and at P12, a majority of genes were those controlling immune response such as *Il4*, *Il6*, *Il2*, and *Tnf* (Fig. 4E, F).

**Figure 4.**
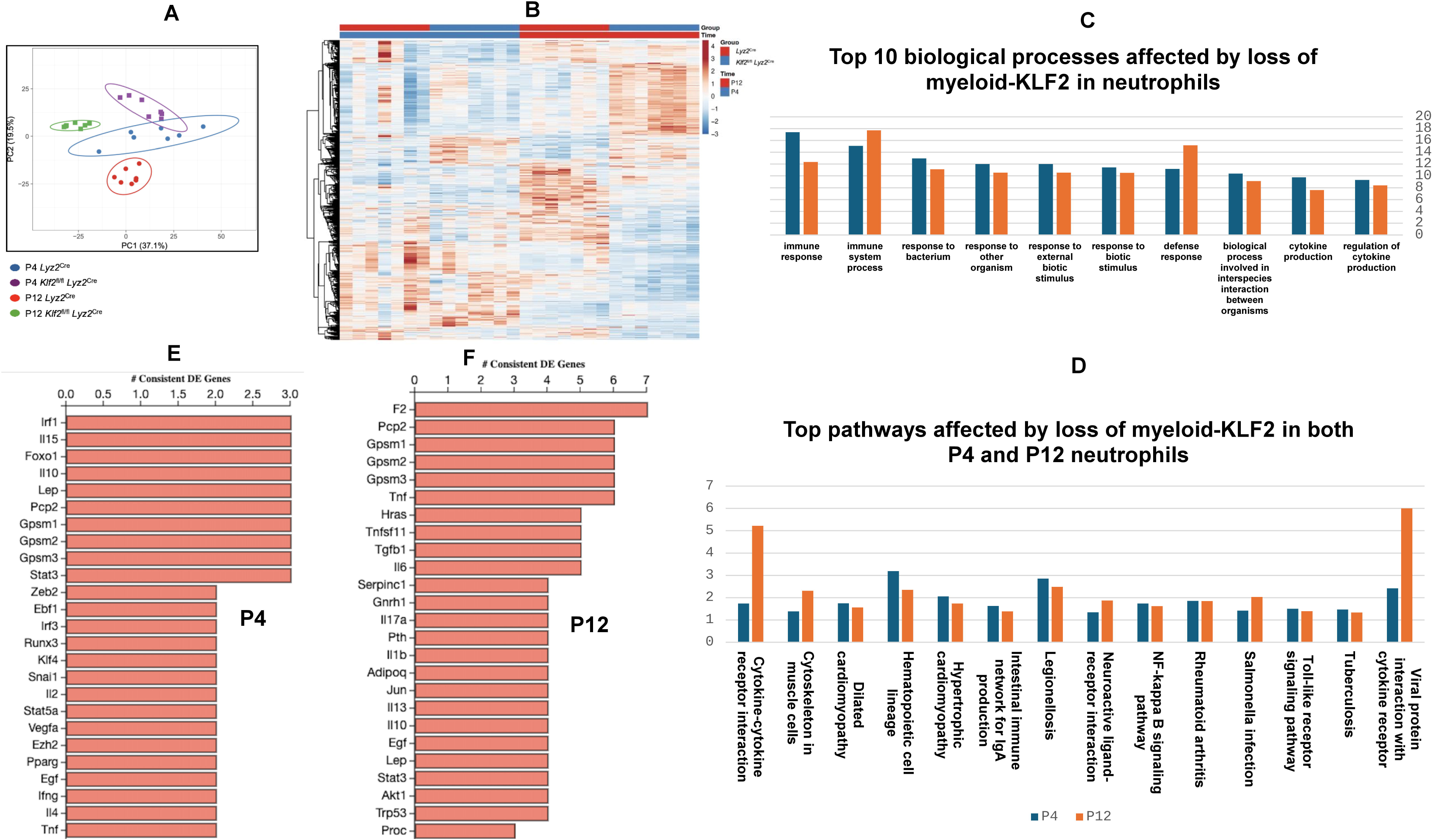
RNA-sequencing of BMNs at P4 and P12 shows enriched inflammatory pathways affected by loss of myeloid-KLF2. (A) Principal component analysis shows separation between the enriched genes between *Klf2*^fl/fl^*Lyz2*^Cre^ and *Lyz2*^Cre^ BMNs at P12 vs. at P4. (B) Heat map showing the topmost differentially expressed genes between 4 groups, n=7 for each group. Individual gene annotations were not made in this heat map due to the number of differentially expressed genes (DEGs) shown in the heat map (C) and (D) show the top biological processes and pathways, respectively, based on a meta-analysis of significant DEGs at P4 and P12 affected by the loss of myeloid-KLF2 from BMNs. Y-axis corresponds to the number of genes present in the pathway or the biological process based on gene ontology analysis enriched at P4 or at P12, between *Klf2*^fl/fl^*Lyz2*^Cre^ and *Lyz2*^Cre^ BMNs. (E) and (F) show the top 25 upstream regulatory genes with targets in the significant DEGs at P4 and at P12. The X-axis indicates the number of downstream targets in the DEGs of the upstream gene, which is annotated on the y-axis.

There were also four distinct upstream regulators in the DEG meta-analysis with consistently differentially expressed targets at both P4 and P12 - *Bach1, Irf3, Mylk*, and *Rfx7*, although these were not in the top 25 upstream regulators at P4 and P12 separately. We then cross-referenced the DEGs in our dataset, which were enriched in immune signature datasets published previously and deposited in the NCBI Gene Expression Omnibus repository separately for P4 and P12. We then selected datasets that incorporated myeloid cells, responses to bacterial antigens, LPS-stimulated cells, and cytokine and chemokine signaling to create a map of the published datasets to the genes enriched between *Klf2*^fl/fl^ *Lyz2*^Cre^ and *Lyz2*^Cre^ at P4 and P12 (Fig. S7, 8). The number of published immune pathway datasets enriched in the significant DEGs in our dataset and their corresponding DEGs in our list of DEGs were more at P12 vs. at P4 (Fig. S7, 8).

In summary, loss of myeloid-KLF2 significantly enriches pro-inflammatory pathways in BMNs by changing the gene expression profile similar to that seen in other inflammatory disease processes, and a lower neutrophil-*Klf2* gene expression at P4 shifts the transcriptome to an pro-inflammatory one at this postnatal age.

## DISCUSSION

Our data are the first to show that neutrophil-KLF2 is a critical regulator of neonatal endotoxemia-related mortality. Others have previously shown that myeloid-*Klf2* deletion leads to increased LPS-induced mortality in 8-week-old adult mice following a four-fold higher dose of LPS than used in our study. Mahabaleshwar et al. used a 21ug/gm LPS dose and identified the macrophage compartment and the HIF-1α pathway as primary drivers of mortality in their adult animal model.[24] This lethal dose causes 100% mortality at P4 and P12 in both *Klf2*^fl/fl^*Lyz2*^Cre^ and *Lyz2*^Cre^ pups (data not shown due to 100% mortality in all arms). By decreasing the dose of LPS to 5ug/gm (for reference, this would be ∼ 10ug for a 2gm P4 pup, 25ug for a 5gm P12 pup, and 125ug for a 25gm adult mouse), we show a developmental influence on the sequelae of endotoxemia, further exacerbated by the loss of myeloid-KLF2. 5ug/gm LPS did not cause mortality in adult mice. Further, a trial of an intermediate dose of 10ug/gm caused 100% mortality at P4 and between 80-100% mortality at P12 (significantly higher mortality in *Klf2*^fl/fl^*Lyz2*^Cre^ vs. *Lyz2*^Cre^ similar to P4, data not shown) to confirm that consequences of endotoxemia are dose, KLF2, and age-dependent. Thus, we chose the 5ug/gm dose for our studies to avoid a 100% mortality rate and enable us to meet various experimental endpoints. Our study sheds new light on the mechanisms operative in the increased mortality observed in preterm neonatal mice compared to term neonatal or adult mice. We demonstrated that neutrophil depletion and not macrophage depletion decreases mortality in the P4 pups, underscoring the critical role of neutrophils in the sequelae of endotoxemia. The observation that a significantly lower *Klf2* gene expression in P4 *Klf2*^fl/fl^ *Lyz2*^Cre^ BMNs is associated with neutrophil-dependent increased mortality at P4 underscores the importance of neutrophil-KLF2 in this process. This is confirmed as myeloid *Klf2*-overexpressing pups have significantly increased survival compared to *Klf2*^fl/fl^ *Lyz2*^Cre^ pups after LPS at P4. We acknowledge that *Klf2* overexpression does not seem to reduce mortality further compared to the control phenotype of *Lyz2*^Cre^, but it is not evident whether this is because of a smaller sample size in the *Klf2*^Tg^ genotype. However, in line with this observation, we did not observe a significant difference in NLRP3 expression or IL-1β release between P4 *Lyz2*^Cre^ and *Klf2*^Tg^ BMNs, although this difference exists between P4 *Klf2*^fl/fl^ *Lyz2*^Cre^ and *Klf2*^Tg^ BMNs. We did not measure *Klf2* gene expression in macrophages between P4 and P12 *Lyz2*^Cre^ pups as macrophage depletion did not decrease mortality.

*Klf2*^fl/fl^*Lyz2*^Cre^ pups have higher serum pro-inflammatory cytokine levels than *Lyz2*^Cre^ pups, particularly pronounced at P4, reflecting the increased mortality at P4. The difference in pro-inflammatory cytokine levels, such as IFN-γ and TNF-α, between *Klf2*^fl/fl^*Lyz2*^Cre^ and *Lyz2*^Cre^ P4 pups were similar to those found previously in adult mice by our group. However, IL-1β was the key pro-inflammatory cytokine whose P4 levels were significantly higher in *Klf2*^fl/fl^*Lyz2*^Cre^ vs. *Lyz2*^Cre^ mice than in the prior adult studies. In addition, IL-1β is elevated in human and animal models of neonatal sepsis.[40–43] IL-18, also produced by NLRP3 activation, is also involved in neonatal sepsis.[44] This adds more evidence that activation of the NLRP3 inflammasome machinery is developmentally dependent and plays a role in neonatal sepsis. We are the first to show that inhibition of NLRP3 inflammasome activation decreases neonatal mortality from endotoxemia.

IL-10, an anti-inflammatory cytokine, was also increased at P4 in *Klf2*^fl/fl^*Lyz2*^Cre^ vs. *Lyz2*^Cre^ pups. Although this mechanism is not fully understood, we outline several potential etiologies for this phenomenon. Firstly, this could be in response to an overwhelming pro-inflammatory cytokine surge in the P4 *Klf2*^fl/fl^*Lyz2*^Cre^ pups. IL-10 is an anti-inflammatory cytokine that protects the body from an uncontrolled inflammatory response.[45] Further, IL-10 release in response to LPS in mice has been shown to increase bimodally, peaking at 1.5h, with a nadir around 6h, and then another increase around 8-12h.[46] We obtained sera at four h after LPS, and the rate of decrease of IL-10 might differ between *Klf2*^fl/fl^*Lyz2*^Cre^ and *Lyz2*^Cre^ pups. Finally, IL-10 release is increased by the presence of apoptotic neutrophils, which is another possibility in *Klf2*^fl/fl^*Lyz2*^Cre^ pups.[47] Our group will explore each of these angles in future studies.

Under physiologic conditions, KLF2 inhibits the recruitment of co-activators p300 and PCAF to NF-κB-binding promoter sites of target genes, including *Nlrp3*.[38, 48] Although both NLRP3 priming and activation were increased in P4 vs. P12 BMNs and further compounded by the loss of myeloid-KLF2, we did not see a significant difference in *Nlrp3* mRNA expression between the four groups based on RNA-seq results, verified by RT-PCR. We acknowledge that we lack data on *Nlrp3* transcript levels in LPS-primed BMNs from the four groups; however, even in the absence of LPS priming, NLRP3 protein expression is higher at P4 and in *Klf2*^fl/fl^*Lyz2*^Cre^ BMNs without any change in *Nlrp3* gene expression.

These data suggest that altered NLRP3 protein levels noted with KLF2 decline are not fully transcription-dependent and may occur through other mechanisms, such as increased phosphorylation of the pyrin or LRR domain, post-translational modifications, or decreased protein degradation. TLR-dependent phosphorylation of the pyrin domain prepares NLRP3 for the priming stimuli.[49] Phosphorylation of serine 803 of the LRR domain is critical in NLRP3 priming, activation, and ubiquitination in murine macrophages, and *Nlrp3*^S803D/S803D^ knock-in mice are resistant to endotoxic shock.[50] Ubiquitination is critical in maintaining NLRP3 at a low level in inactivated cells, primarily by K48-linked Ub chain-dependent proteasomal degradation and K63-linked Ub chain-dependent autophagic degradation.[51–53] An investigation into the role of KLF2 in these mechanisms is underway.

*Klf2*^fl/fl^ *Lyz2*^Cre^ pups have increased pro-IL-1β cleavage in their LPS-primed BMNs and release of mature IL-1β and caspase-1 p20 into the supernatant, demonstrating that NLRP3 activation is heightened in these BMNs. We cannot conclude whether the increased cleavage of pro-IL-1β and pro-caspase-1 are entirely attributable to increased NLRP3 priming seen in *Klf2*^fl/fl^ *Lyz2*^Cre^ BMNs or whether there is a separate increase in the activation step (signal 2). Assembly of the NLRP3 inflammasome involves the interaction of the NLRP3 protein with the N-terminus of an adaptor protein, ASC, via pyrin-pyrin interactions.[54] The C-terminus of ASC has a caspase recruitment domain (CARD) that can bind to pro-caspase-1 through CARD-CARD interactions, leading to the dimerization of pro-caspase-1 molecules.[55] Caspase-1 dimerization leads to activation and proteolytic cleavage of pro-caspase-1 into caspase-1 p33/p10, which cleaves pro-IL-1β into mature IL-1β and subsequently self-cleavage into p20 and p10 species, which are released into the supernatant.[55] The observation that unprimed P4 *Klf2*^fl/fl^ *Lyz2*^Cre^ BMNs have caspase-1 p20 release into the supernatant does indicate that there is increased NLRP3 assembly even in the absence of priming in *Klf2*^fl/fl^ *Lyz2*^Cre^ BMNs. It is also possible that non-NLRP3 inflammasomes, such as AIM2, which can trigger caspase-1 cleavage without needing a prior TLR-mediated priming stimulus, unlike the NLRP3 inflammasome, are contributing to this phenomenon.[56] This observed phenomenon might explain why unprimed P4 *Klf2*^fl/fl^ *Lyz2*^Cre^ BMNs have no expression of pro-IL-1β in their whole cell lysates, and there is a minute but detectable expression of mature IL-1β in the supernatant in these cells, likely due to proteolytic cleavage by caspase-1 p20. We do not observe a difference in *Aim2* transcript levels in our transcriptomic analysis, but further work into these mechanisms is underway.

Importantly, we found no significant difference in *Hif-1α* mRNA levels between the four arms in our RNA-seq results, unlike adult *Klf2*^fl/fl^*Lyz2*^Cre^ mice subjected to endotoxemia. We demonstrate that neutrophil-NLRP3 is critical to endotoxemia-mediated mortality in neonates. We acknowledge that MCC950 globally inhibits NLRP3 activation in vivo and is not specific to BMNs. However, our data that (1) neonatal LPS-mortality is neutrophil-dependent, (2) neutrophil-specific NLRP3 and IL-1β release are higher in animals experiencing higher mortality, and (3) NLRP3 inhibition decreases mortality strongly suggest that the increased mortality in neonatal pups is at least partly dependent on neutrophil-NLRP3 priming and activation.

The more significant decrease in *Klf2* gene expression at P4 vs. at P12 between *Klf2*^fl/fl^ *Lyz2*^Cre^ vs. Lyz2^Cre^ neutrophils (Fig. S6) likely contributes to the stronger phenotype in the *Klf2*^fl/fl^ *Lyz2*^Cre^ pups at P4 vs. at P12. The *Klf2*^fl/fl^ *Lyz2*^Cre^ strain constitutively expresses the Cre recombinase protein and is homozygous for the floxed-*Klf2* gene. However, postnatal changes in upstream regulatory genes could contribute to the compensatory increase in *Klf2* gene expression from P4 to P12 in the *Klf2*^fl/fl^ *Lyz2*^Cre^ strain, which still has a lower *Klf2* gene expression than the *Lyz2*^Cre^ strain at P12 (Fig. S6). Further work by our group on the mechanism underlying the influence of postnatal age on *Klf2* gene expression in neutrophils in different mouse strains is ongoing. These studies will identify the etiology contributing to increased mortality from endotoxemia at earlier postnatal ages, regardless of genetic composition. Of the four upstream regulators identified in the pathway analysis, *Bach1* and *Irf3* are the only genes associated with NLRP3 signaling. Studies in murine *Bach1*-knockout macrophages demonstrate increased NLRP3 activation in response to LPS and nigericin through enhanced generation of mitochondrial reactive oxygen species.[57] As NLRP3 transcript levels are not altered by Bach1, this could be a possible mechanism to explain our results. However, this does not explain the increased NLRP3 protein expression in P4 *Klf2*^fl/fl^*Lyz2*^Cre^ BMNs. Type-1 interferon-regulatory factor-3 (IRF3) is a transcription factor downstream of the *Trif* gene (NF-kB regulator) and increases gene expression of type-1 interferons, which affects NLRP3 activation.[58–60] Stimulator of interferon genes (STING) binds to IRF3 and phosphorylated IRF3, which translocates to the nucleus and increases the expression of *Nlrp3*.[60] IRF3 is less likely to be the sole explanation of our results because there are no differences in NLRP3 transcript levels between the four groups. Our future investigations will focus on the specific cause of increased NLRP3 priming from the loss of myeloid-KLF2 at P4, in addition to the mechanism of increased caspase-1 cleavage induced by loss of KLF2 in BMNs, and the contribution of these phenomena to KLF2-mediated changes in the neonatal innate immune response to bacterial antigens.

Our studies show three critical findings: (i) neutrophils are key players in the increase in mortality noted at an earlier postnatal age; (ii) *Klf2* gene expression is associated with increased NLRP3 priming and activation in a postnatally-dependent fashion; (iii) NLRP3 priming and activation contributes significantly to neonatal mortality from endotoxemia as noted by survival benefit following blockade of NLRP3 activation. Recent studies have challenged the notion that the neonatal immune system is shifted towards a more anti-inflammatory response, and emerging evidence points towards a pro-inflammatory phenotype. Neonatal mice are paradoxically hyper-responsive to LPS and other TLR agonists. After LPS stimulation, the increase in pro-inflammatory cells in preterm neonates is more pronounced and faster than in adults.[61, 62] The compensatory anti-inflammatory response system in preterm neonates is immature, with profoundly decreased IL-10 production.[63–65] Most neonatal studies on sepsis primarily focus on the adaptive immune system or the monocyte/macrophage compartment. Our analysis shows the critical role of neutrophils in regulating the response to endotoxemia in neonates and that within the neutrophil compartment, KLF2 and NLRP3 are important determinants of increased sepsis-related mortality, a hallmark of this particular age group. KLF2 maintains myeloid cells in a quiescent state, and a considerably lower *Klf2* expression in P4 neutrophils potentially contributes to the pro-inflammatory cytokine storm-driven mortality seen in P4 *Klf2*^fl/fl^ *Lyz2*^Cre^ pups. Our novel findings form the basis for future studies targeted towards NLRP3 inhibition and KLF2-targeted therapies for neonatal sepsis, which to this day remains associated with a high mortality burden.

## Acknowledgments

We acknowledge Dr. Tracey Bonfield (CWRU) for allowing the generous use of the FLIR thermal imager. The Genomics Core Facility of the CWRU School of Medicine supported this research. The featured image was Created in BioRender, *Mukherjee, D. (2025)* https://BioRender.com/p69l853. This work was supported in supplies and effort by NIH R01HL142647 (LN), NIH P30CA043703 (AYH), Hyundai Hope-on-Wheels Scholar Award (AYH), CURE Childhood Cancer Translation to Cure Award (AYH), NIH P01AI41350 (GRD), Core Utilization Pilot (DM) from CTSC of Cleveland (NCATS UL1TR0002548), Rainbow Babies Faculty Pilot Award (DM), and Neonatal Research Grant from the Little Giraffe Foundation (DM).

## Authorship and conflicts of interest

The authors contributed to the manuscript as stated below: Study conception and design: DM and LN. Execution of experiments: DM, SS, AT, YL, SC, KW. Analysis and interpretation of results: DM, SS, AT, ERC, LN. Draft manuscript preparation: DM, AT, AYH, MKJ, GD, LN. All authors reviewed the data in the manuscript and approved the final version of the manuscript.

All authors declare no competing financial interests.

## Data Availability

The sequencing dataset, with the accession code GSE278604, is available from the NCBI GEO repository. Upon a reasonable request, the corresponding author will provide all original data.

