## Supplementary Data for "Neutrophil KLF2 regulates inflammasome-dependent neonatal mortality from endotoxemia"

Table S1. Real-time PCR primer sequences.

| Gene and template direction | 5’->3’ sequence |
| --- | --- |
| KLF2 Forward | CACCTAAAGGCGCATCTGCGTA |
| KLF2 Reverse | GTGACCTGTGTGCTTTCGGTAG |
| NLRP3 Forward | TCACAACTCGCCCAAGGAGGAA |
| NLRP3 Reverse | AAGAGACCACGGCAGAAGCTAG |
| RPL13A Forward | ATGACAAGAAAAAGCGGATG |
| RPL13A Reverse | CTTTTCTGCCTGTTTCCGTA |

Table S2. Hallmark signature pathways that were significantly enriched in the RNA-seq dataset at P12.

| Hallmark pathway | Number of genes enriched | P value |
| --- | --- | --- |
| TNF-a signaling via NF-kB | 29/195 | <0.0001 |
| Inflammatory response | 29/196 | <0.0001 |
| Coagulation | 17/132 | 0.001 |
| Kras signaling | 19/192 | 0.011 |
| IL-6 JAK STAT3 Signaling | 10/83 | 0.036 |
| IL-2 STAT5 signaling | 17/192 | 0.043 |
| Myogenesis | 17/196 | 0.046 |

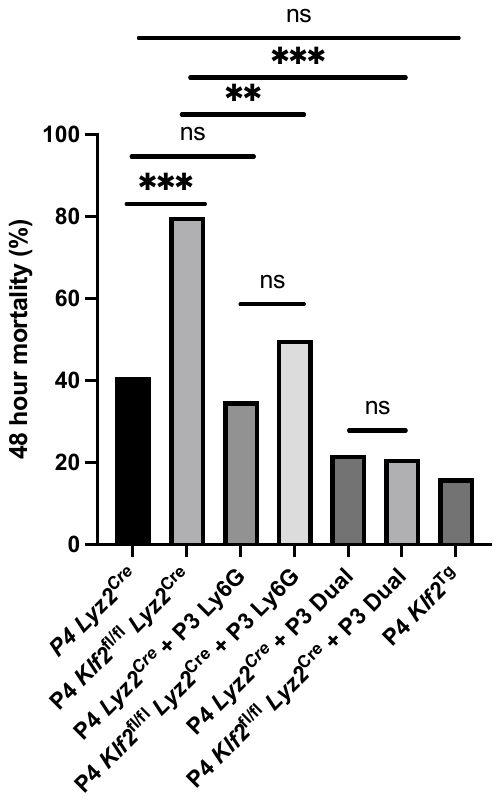

Figure S1. 48-hour mortality after 5ug/gm intraperitoneal LPS injection in P4 pups. P3 Ly6G regimen consisted of 100ug/gm anti-Ly6G 1A8 clone injection 24 hours before LPS injection, and P1-3 Ly6G regimen consisted of daily anti-Ly6G 1A8 clone injections from P1 – P3, followed by LPS injection on P4. For LPS-only studies, *Klf2*^fl/fl^ *Lyz2*^Cre^ pups had an n=40, *Lyz2*^Cre^ pups had an n=39, and *Klf2*^Tg^ had an n=19. For all antibody depletion studies, the number of animals was 30 in each group. ** = p<0.01, *** = p<0.001, and ns= not significant.

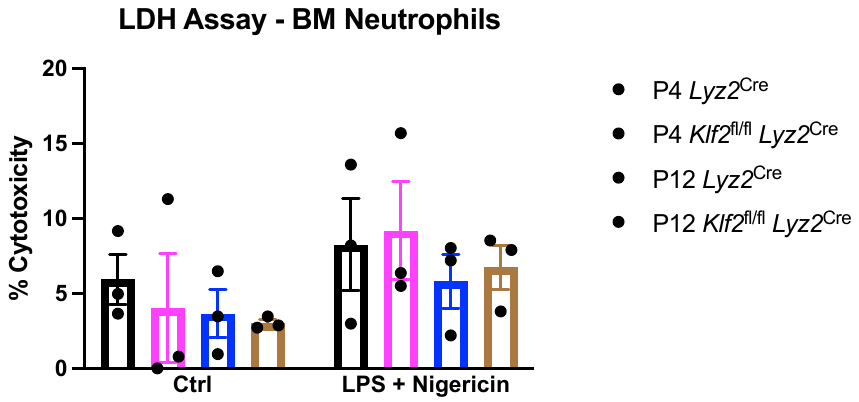

ns

Figure S2. Lactate dehydrogenase assay to detect cytotoxicity- Bone marrow neutrophils were isolated and plated at 10,000 cells/100uL seeding density in 96-well plates in RPMI-1640 media, and treated with LPS for 2 hours, followed by nigericin for 30 mins. BMNs were also treated with lysis buffer in separate wells for the whole duration toi detect maximal LDH release. The supernatant was aspirated and used to detect LDH activity using fluorescent plate reader. Cytotoxicity was calculated as the percentage of the maximal LDH activity for each cell type and treatment arm. LPS priming followed by nigericin stimulation leads to increased cytotoxicity, but this increase was not statistically significant across any of the four groups (two-way ANOVA). Each data point represents individual biological replicates.

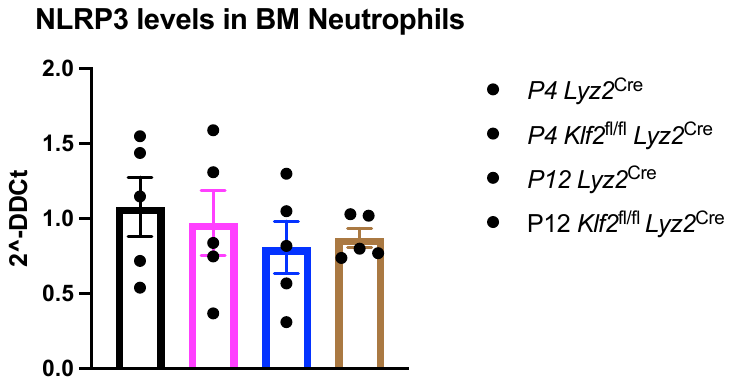

Figure S3. Unstimulated BMNs were used for RT-PCR to quantify NLRP3 expression, and there was no significant difference in NLRP3 mRNA transcript levels between the four arms. Ct values were normalized to RPL13A to obtain ΔCt, followed by normalization to P4 *Lyz2*^Cre^ to obtain ΔΔCt values. N=5 for each group.

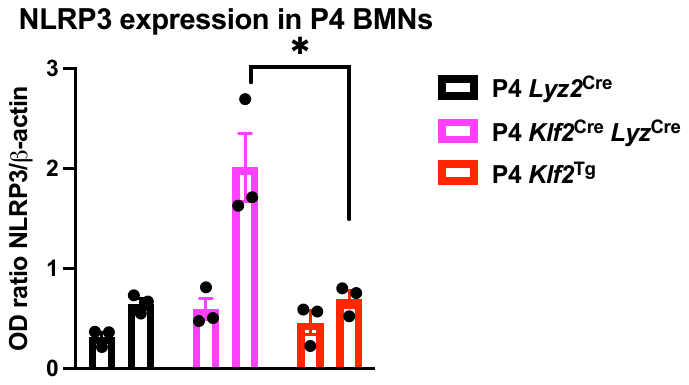

Figure S4. Protein expression of NLRP3 (117kDa) normalized to beta-actin in P4 *Klf2*^fl/fl^ *Lyz2*^Cre^, *Lyz2*^Cre^, and *Klf2*^Tg^ BMNs at baseline and primed and activated with LPS and nigericin. Primed and activated *Klf2*^Tg^ BMNs have significantly lower NLRP3 expression compared to *Klf2*^fl/fl^ *Lyz2*^Cre^ BMNs (p=0.02). P4 *Klf2*^fl/fl^ *Lyz2*^Cre^ and *Lyz2*^Cre^ were run on the same gel, and *Klf2*^Tg^ was run on a separate gel but under the same conditions (4-20% sodium dodecyl sulfate-polyacrylamide gel electrophoresis at 120V) and with similar protein loading amounts, and normalized to beta-actin.

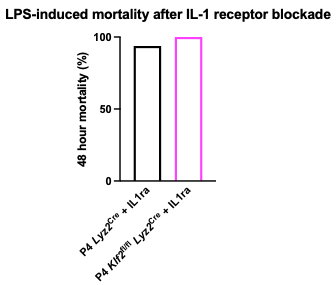

Figure S5. 48-hour mortality after 2 days of IL-1 receptor antibody (BioXCell #BE0256) on P3 and P4, 24 hours apart, followed by 5ug/gm intraperitoneal LPS injection on P4, 4 hours after the last antibody injection. *Klf2*^fl/fl^ *Lyz2*^Cre^ pups had an n=17, *Lyz2*^Cre^ pups had an n=16. P4 *Klf2*^fl/fl^ *Lyz2*^Cre^ pups had 100% survival after IL-1 receptor blockade, whereas P4 *Lyz2*^Cre^ pups had a 93.8% survival (15/16), 48 hours after LPS.

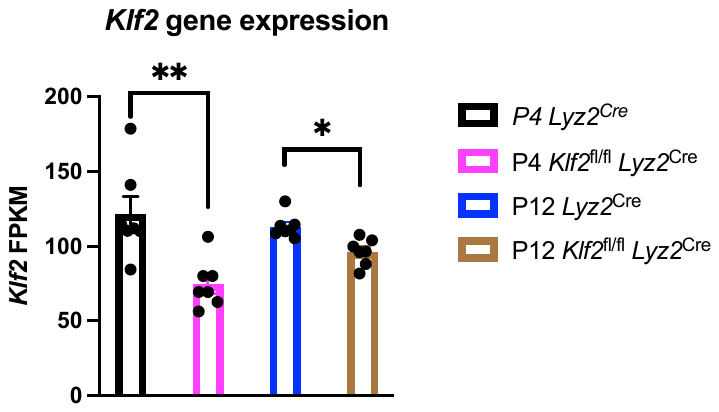

Figure S6. *Klf2* gene expression from transcriptomic analysis is expressed as Fragments Per Kilobase of transcript per Million mapped reads (FPKM) on the left y-axis.

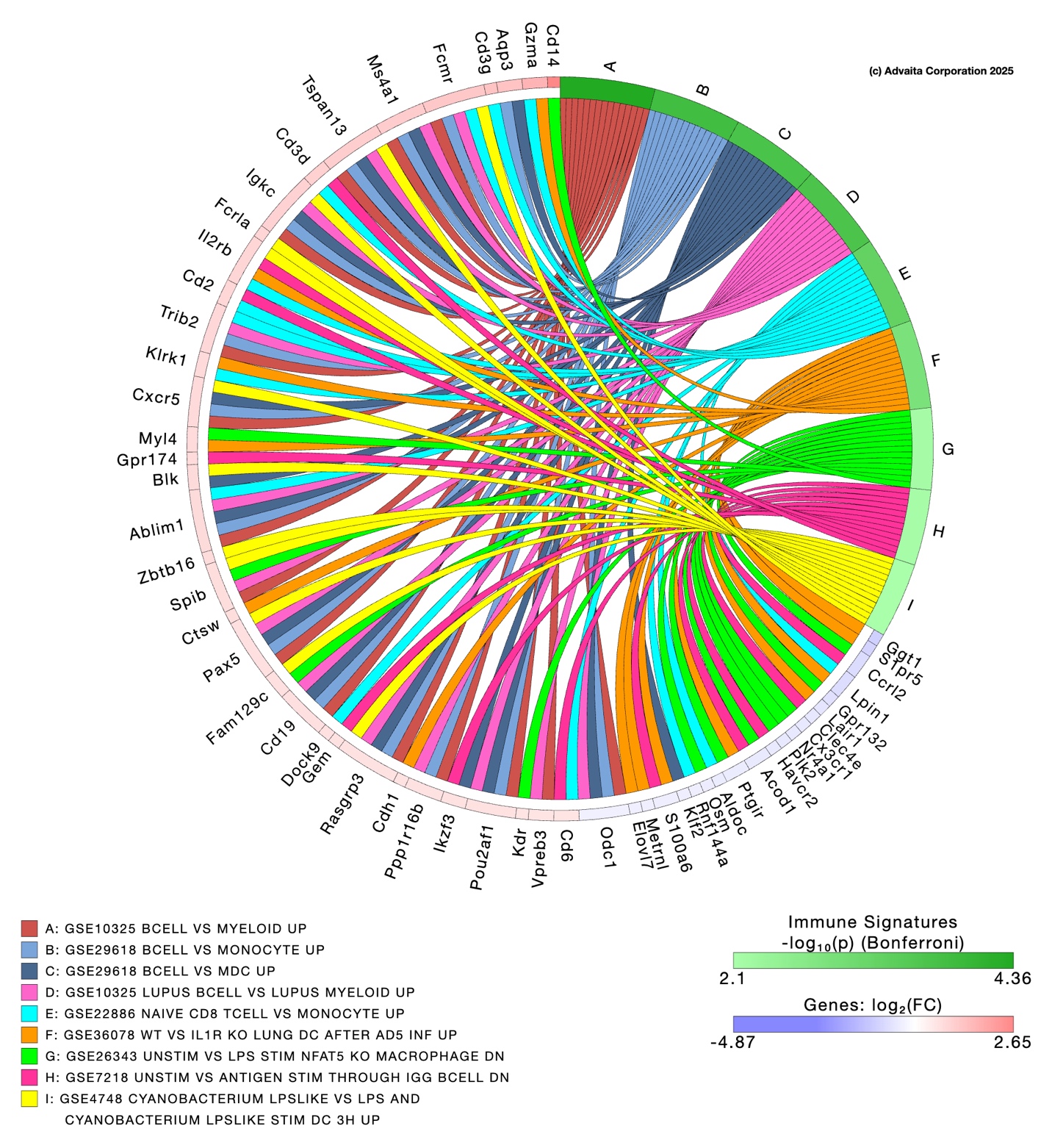

Figure S7. Using iPathway Guide, significant DEGs at P4 were cross-referenced to published RNA-sequencing datasets with the inbuilt immune signature tool. Datasets with relevance to our study, including myeloid cell response, effect of LPS stimulation, cytokine and chemokine signaling, etc. were included to identify the DEGs in our dataset which were enriched in these published datasets. All GEO accession numbers of individual datasets are listed in the key on the left bottom corner. Green-shaded circular ring to the immediate outside of the datasets refer to the level of significance with which each dataset had enriched genes in our dataset, and blue-to-red shaded ring to the immediate outside of the annotated genes refer to the fold change of each gene in our dataset.

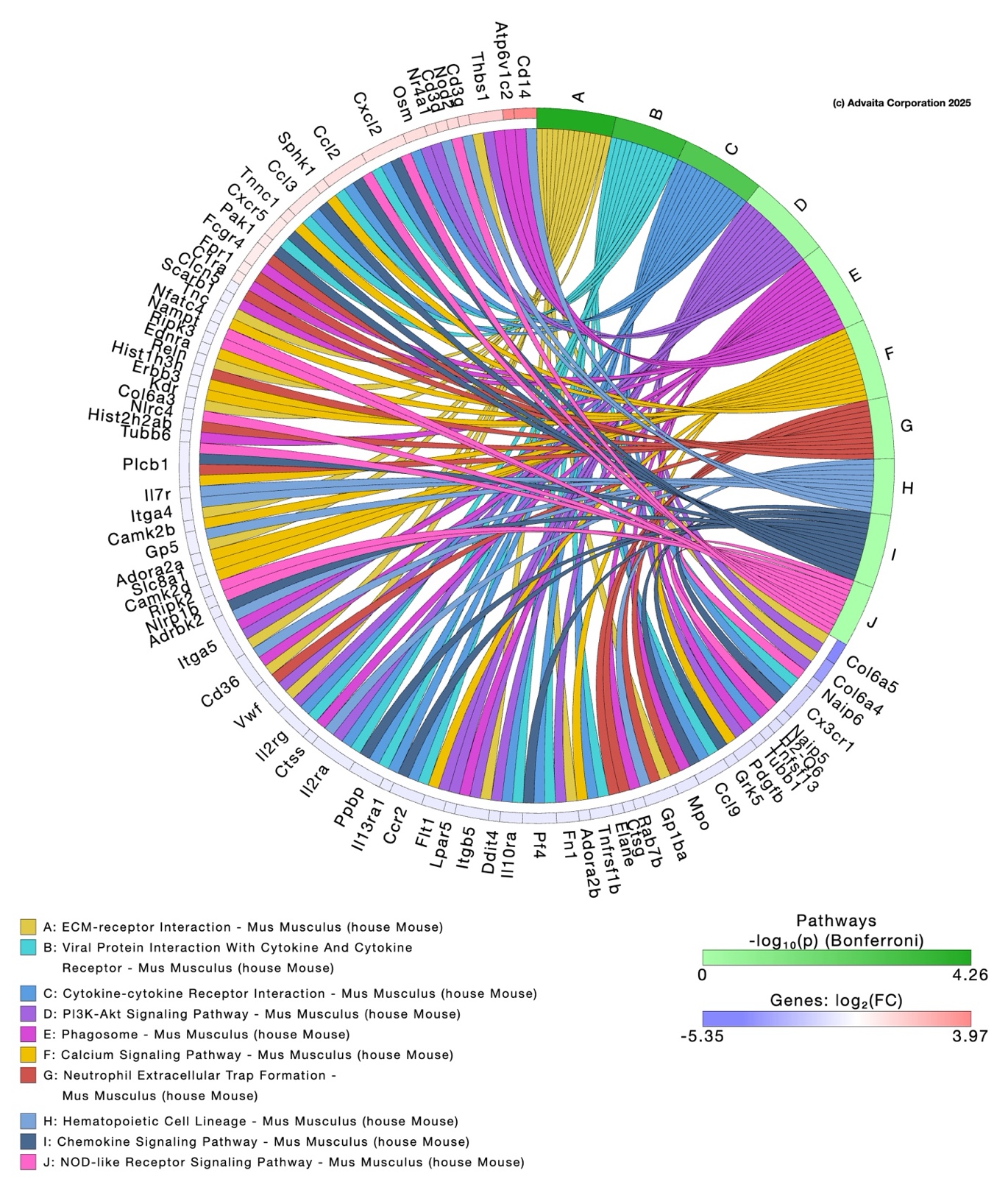

Figure S8. Figure S7. Using iPathway Guide, significant DEGs at P12 were cross-referenced to published RNA-sequencing datasets with the inbuilt immune signature tool. Datasets with relevance to our study, including myeloid cell response, effect of LPS stimulation, cytokine and chemokine signaling, etc. were included to identify the DEGs in our dataset which were enriched in these published datasets. All GEO accession numbers of individual datasets are listed in the key on the left bottom corner. Green-shaded circular ring to the immediate outside of the datasets refer to the level of significance with which each dataset had enriched genes in our dataset, and blue-to-red shaded ring to the immediate outside of the annotated genes refer to the fold change of each gene in our dataset.
